# Radiation-induced disruption of cardiac mitochondrial bioenergetics and nucleotide homeostasis in mice

**DOI:** 10.64898/2026.06.08.730816

**Authors:** Klaudia Stawarska, Ada Kawecka, Krzysztof Urbanowicz, Joanna Kamińska, Michał Posiewnik, Alicja Braczko, Wiktoria Michnowska, Barbara Kutryb-Zając, Bartłomiej Tomasik

## Abstract

**Aims:** Cardiac stereotactic body radiotherapy (SBRT) has emerged as a promising non-invasive treatment for refractory ventricular tachycardia (VT). Intriguingly, the clinical benefit of SBRT often occurs within days of treatment, preceding the development of radiation-induced fibrosis, suggesting alternative underlying mechanisms. This study aimed to investigate the acute and persistent effects of ionizing radiation on cardiac bioenergetics and mitochondrial function, providing mechanistic insights into early cardiac responses to radiation exposure.

**Methods and results:** We employed a translational multi-model approach, including HL-1 mouse cardiomyocytes and *ex vivo* mouse left ventricular living myocardial slices (LMS). Bioenergetic profiling, assessment of mitochondrial respiration and calcium handling were performed following exposure to clinically relevant radiation doses (10 Gy and 25 Gy). In HL-1 cardiomyocytes, 10 Gy induced acute bioenergetic stress, characterized by reduced adenylate energy charge, cytoskeletal disorganization, and impaired mitochondrial respiration, accompanied by increased calcium oscillation amplitude. 25 Gy exposure led to NAD^+^ depletion but paradoxically enhanced mitochondrial respiratory capacity, suggesting an adaptive metabolic response. Murine myocardial slices demonstrated reduced creatine content while preserving energy balance as indicated by phosphocreatine/ATP ratio, indicating tissue-level metabolic resilience. These findings reveal model-specific metabolic perturbations induced by cardiac irradiation, underscoring the importance of tissue complexity in modulating the cardiac response to radiation.

**Conclusion:** This study demonstrates that ionizing radiation at 10 Gy and 25 Gy induced dose- and model-dependent bioenergetic alterations in cardiac cells and tissues, including changes in mitochondrial respiration, nucleotide levels, and redox balance. While 10 Gy exacerbated metabolic disruption, 25 Gy triggered partial recovery, highlighting differential responses across cellular and tissue levels. These metabolic changes may contribute to the immediate effects of cardiac SBRT and potentially to long-term cardiotoxicity.

**Translational Perspective:** Our study provides novel mechanistic insights into the metabolic effects of cardiac irradiation, revealing acute mitochondrial stress, redox imbalance and alterations in calcium homeostasis in cardiomyocytes. These early bioenergetic changes may contribute to both the immediate anti-arrhythmic effects and the potential long-term cardiotoxicity of stereotactic body radiation therapy. Understanding these molecular responses is essential to optimize the therapeutic window of cardiac radioablation and minimize adverse effects.

**GRAPHICAL ABSTRACT:** 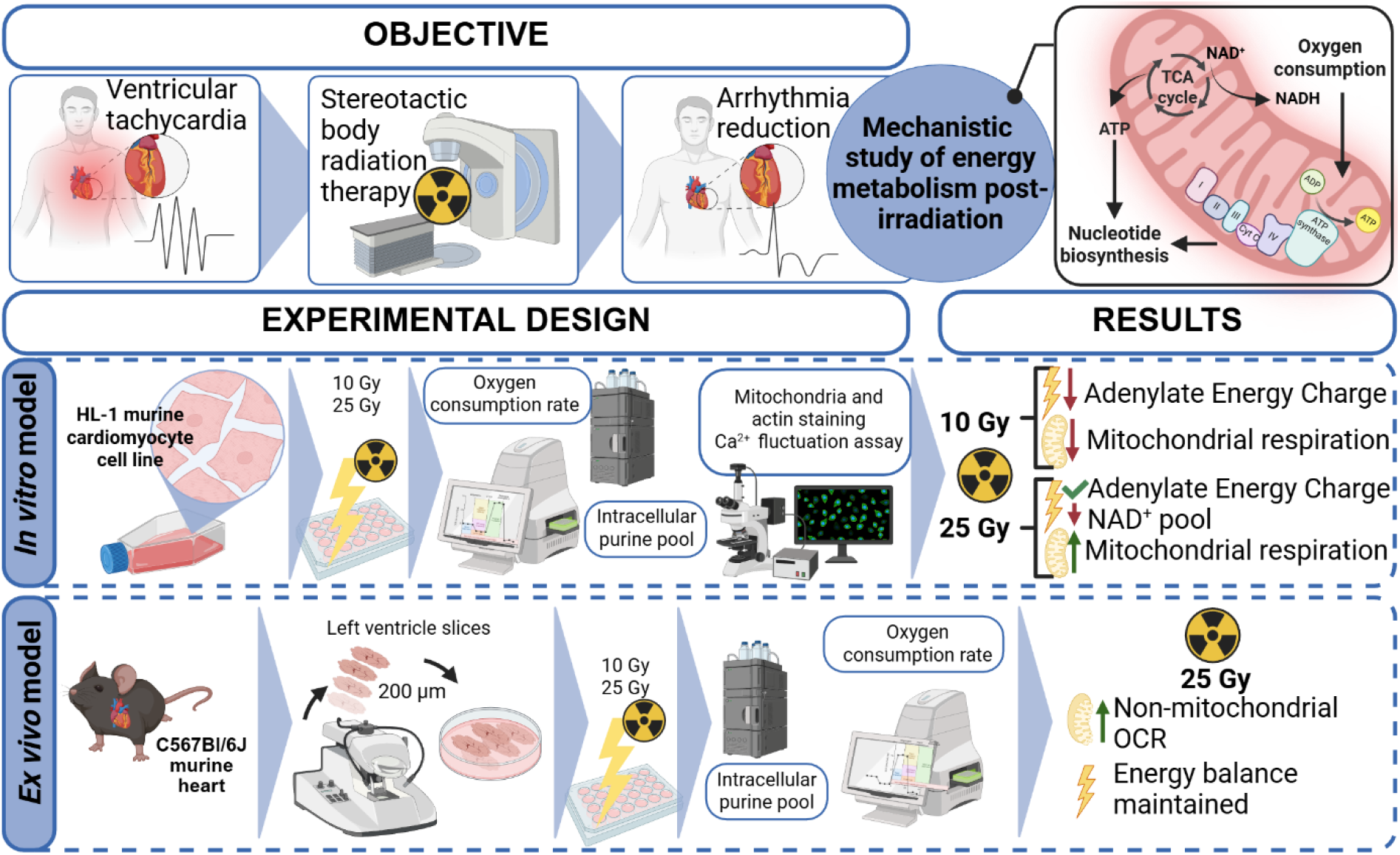

## INTRODUCTION

Stereotactic body radiation therapy (SBRT) has emerged as an effective non-invasive treatment option for life-threatening ventricular tachycardia (VT), particularly in patients refractory to standard interventions such as anti-arrhythmic medications or catheter ablation ^1,2^. A single high-dose irradiation, typically 25 Gy, delivered in an outpatient setting to the limited volume of the heart without the need for anaesthesia, has been associated with significant reductions in arrhythmia burden and ICD interventions^3^. Several clinical studies have demonstrated the rapid and profound anti-arrhythmic effects of SBRT, with a significant reduction in arrhythmia burden often observed within days to weeks after treatment^4–7^. This early clinical benefit precedes the development of radiation-induced fibrosis, which traditionally has been considered the primary mechanism underlying the therapeutic effect of cardiac irradiation^8,9^.

The underlying biological mechanisms mediating the immediate and sustained effects of cardiac irradiation remain incompletely understood. While initial hypotheses emphasized radiation-induced scar formation and structural conduction block, recent preclinical and clinical evidence indicates that functional electrophysiological and metabolic reprogramming of cardiomyocytes may play a critical role^10^. Experimental data suggest that ionizing radiation may exert direct effects on cardiomyocytes by modulating their electrophysiological properties. Preclinical studies have demonstrated acute alterations in the expression of ion channels, calcium-handling proteins, and gap junction components following irradiation, leading to increased conduction velocity, shortened action potential duration, and enhanced cell-to-cell coupling^11–13^. Emerging evidence indicates that radiation-induced oxidative stress and microvascular injury may contribute to arrhythmia suppression and tissue remodelling ^14,15^. Ionizing radiation creates reactive oxygen species (ROS) directly *via* water radiolysis, which in turn interact with mitochondrial membrane lipids, causing disruption of mitochondrial membrane integrity and further increasing ROS level intracellularly. The resulting redox imbalance drives degradation of nucleic acids, lipids and proteins, ultimately leading to necrotic cell death^16^. Recent findings also show that whole-heart irradiation of mice with an endothelial cell-specific p53 deletion, delivered as a single 12 Gy dose, induces increased myocardial vascular permeability, reduced vascular density, and enhanced cell death^17^.

However, beyond these electrophysiological and structural changes, there is growing recognition of the critical role of mitochondrial dysfunction and metabolic reprogramming in the pathophysiology of radiation-induced cardiac injury^18^. Cardiomyocytes are highly energy-dependent cells, relying primarily on mitochondrial oxidative phosphorylation to sustain their contractile and electrophysiological activity^19^. Disruption of mitochondrial function and bioenergetics can compromise cellular calcium homeostasis, membrane excitability, and overall cardiac performance. Ionizing radiation is known to impair mitochondrial integrity, increase ROS production, and induce changes in nucleotide metabolism, yet the impact of therapeutic radiation doses on cardiac metabolic pathways remains insufficiently characterized^20–22^.

To date, few studies have systematically addressed how ionizing radiation affects cardiac bioenergetic homeostasis and whether these metabolic alterations contribute to the observed anti-arrhythmic effects or, conversely, to long-term cardiotoxicity. In particular, the dynamics of intracellular nucleotide pools, adenylate and guanylate energy charge, and nicotinamide adenine dinucleotide (NAD^+^/NADH) balance in response to irradiation have not been comprehensively explored^10,12,23^. Nevertheless, NAD^+^ depletion during heart failure has been associated with the occurrence of spontaneous lethal arrhythmias, indicating a significant impact on cardiac electrophysiology^24^.

In this study, we aimed to bridge this knowledge gap by investigating the acute effects of ionizing radiation on mitochondrial function and intracellular energy metabolism in cardiac cells and tissues. Using a multi-model approach that included murine HL-1 cardiomyocytes and *ex vivo* left ventricular myocardial slices, we assessed the dose-dependent alterations in nucleotide profiles, mitochondrial function and architecture following exposure to clinically relevant doses of irradiation^25,26^.

## METHODS

### Animal handling and heart retrieval

Six three-month-old male C57BL/6J (Wild-Type; WT) mice were fed water and a standard chow diet. All experiments followed the guidelines for the Care and Use of Laboratory Animals published by the European Parliament, Directive 2010/63/EU. Animals were kept in individually ventilated cages under controlled environmental conditions (55 ± 10% humidity, 23 ± 2 °C) with a 12-hour light/dark cycle. Anaesthesia was administered intraperitoneally with a mixture of ketamine (100 mg/kg) and xylazine (10 mg/kg). Whole hearts were removed from the chest and quickly placed in warm (37 °C) slicing buffer, containing heparin sodium at final concentration 2 IU/ml. After pumping out the blood and atrial cutting, hearts were transferred to cold (4 °C) slicing buffer, where the left ventricle was isolated and prepared for slicing.

### Myocardial slicing and incubation

The left ventricular fragments of mouse hearts were mounted to a specimen holder with tissue adhesive, epicardium down ^27^. Myocardial slicing was carried out in a slicing buffer with pH 7,35, containing 0.9 mM CaCl_2_, 5,4 mM KCl, 136 mM NaCl, 1 mM MgCl2, 0.33 mM NaH2PO4, 10 mM glucose, 10 mM HEPES and 30 mM 2,3-butanedione monoxime (cat. no. B0753, Merck, Germany), continuously oxygenated with carbogen and maintained at a temperature of 4 °C. We used a 7000smz-2 vibratome (Campden Instruments Ltd., Loughborough, Leics., England) and the operating parameters of the microtome were as follows: section thickness - 200 µm, blade speed - 0.03 mm/s, vibration frequency - 80 Hz, amplitude - 2 mm. Isolated slices were rinsed in phosphate-buffered saline (cat. no. 21-040-CV, Corning, Manassas, VA, USA) with 3% penicillin/streptomycin (P/S) (cat.no. 30-002-CI, Corning, Manassas, VA, USA) and incubated for one hour at 37 °C, 5% CO_2_ in medium 199 (cat. no. M2154, Merck, Germany) with 1% ITS (cat. no. I3146, Merck, Germany) and 1% P/S until further irradiation.

### Experimental design

Samples were exposed to low (10 Gy) and high (25 Gy) ionizing radiation. Cardiac slices and HL-1 cells irradiation was done at the Department of Oncology and Radiotherapy, Medical University of Gdańsk, Poland, using a linear accelerator (Varian TrueBeam®) delivering a single 10 × 10 cm flattening filter free beam, with a single dose (10 or 25 Gy). The prescribed doses of 10 Gy and 25 Gy were delivered in a single fraction with a dose rate of 600 monitor units (MU)/minute. The corresponding monitor units were approximately 850 MU for 10 Gy and 2125 MU for 25 Gy, resulting in beam-on times determined automatically by the linear accelerator based on these parameters. Approximately one hour after treatment, slices were preserved in liquid nitrogen and lyophilized in a freeze-dryer immediately. Alternatively, cardiac slices, used as a control group, were transported to the clinic but were not irradiated.

### Intracellular nucleotide measurement

Dried cardiac tissues were homogenized in a glass homogenizer with 0,4M HClO_4_. After 15 minutes incubation on ice and 15 minutes centrifugation at 14000 RPM, the supernatant was neutralized with 2M KOH. The incubation and centrifugation step was then repeated, and samples were analysed with ultra-high performance liquid chromatography (UHPLC), as described earlier ^28^.

### Mitochondria function analysis in cardiac slices

The mitochondrial function of cardiac tissue slices was assessed using the Agilent Seahorse XF Flex analyzer with the Seahorse XF Flex 3D Capture Microplate-L, in accordance with the manufacturer’s instructions. Briefly, cardiac tissue was sectioned into 200 µm thick slices and punched into 2 mm diameter samples to fit the microplate wells. One day prior to the assay, the sensor cartridge was hydrated with 1 mL of Seahorse XF calibrant solution per well and incubated at 37 °C in a non-CO_2_ incubator overnight. On the day of the experiment, samples were loaded into wells containing assay medium, and the capture screen was positioned to secure the tissue. The mitochondrial stress test was conducted by sequential injection of compounds in the following order: oligomycin A at a final well concentration of 30 µM, FCCP at 20 µM, and a mixture of rotenone and antimycin A each at 10 µM. Oxygen consumption rates were recorded to measure mitochondrial respiration parameters including basal respiration, ATP-linked respiration, maximal respiration, proton leak and non-mitochondrial respiration in the 3D cardiac tissue slices.

### Cell culture and treatment

Murine cardiac muscle cell line HL-1 (cat. no. SCC065, Merck, Germany) was cultured in a downright humid atmosphere at 37 °C and 5% CO_2_, in gelatin- and fibronectin-coated flasks (EmbryoMax 0.1% Gelatin Solution cat. no. ES-006, Merck, Germany; fibronectin-F1141, Merck, Germany) in Claycomb Medium (cat. no. 51800C, Merck, Germany) supplemented with 0.1 mM norepinephrine (cat. no. A0937, Merck, Germany), 0.3 mM L-Ascorbic Acid (cat. no. A7506, Merck, Germany), 10% (v/v) FBS (cat. no. 10500064, Gibco, Brazil), 2 mM L-glutamine (cat. no. 25-005-CV, Corning, Manassas, VA, USA) and 1% (v/v) penicillin/streptomycin (cat. no. 30-002-CI, Corning, Manassas, VA, USA). Cells were seeded on coated multi-well plates 48 hours before the treatment. 24 hours before treatment, the cell medium was altered to one with reduced FBS concentration (1%). Cells were exposed to 10 or 25 Gy of radiation; all experimental procedures were conducted approximately 1 hour after treatment.

### HL-1 intracellular nucleotide concentration quantification

Cells were seeded on 24-well plates at 0.05 × 10^6^ cells per well. After the treatment, the cell monolayer was rinsed two times with HBSS and frozen in ice-cold 0.4 M HClO_4_ at -80 °C for 24 hours, thawed, and frozen again. Collected supernatants were centrifuged for 10 minutes at 14 000 RPM, 4 °C, and neutralized to pH 5.5-6.5 with 3M K_3_PO_4_, centrifuged again, and the supernatants were analysed using UHPLC, as previously described ^29^. The residue on the plate was air-dried and dissolved in 0,5M NaOH to determine protein content in each sample using the Bradford method (cat. no. 5000006, Bio-Rad, Hercules, CA, USA).

### Mitochondria function analysis in cells

Mitochondria function analysis was performed using a Seahorse XFp analyzer (Agilent, Santa Clara, CA, USA) ^30^. Cells were seeded on XFp 8-well plates at 0.01 × 10^6^ cells per well. After the exposure to the radiation, cells were washed and incubated in Seahorse DMEM medium supplemented with 1 mM pyruvate, 2 mM glutamine, and 10 mM glucose at 37 °C for 45 minutes. Then, XF Cell Mito Stress Test was performed according to the manufacturer’s instructions. We used commercially available assays to assess mitochondrial respiration, and the concentrations of respiratory chain inhibitors were selected according to the manufacturer’s recommendations and validated for group responsiveness. However, only a single FCCP concentration was applied, and the absence of a full FCCP titration precludes excluding potential group-specific differences in uncoupler sensitivity or mitochondrial membrane potential. Briefly, the analyzer sequentially injected oligomycin (inhibitor of Complex V), carbonyl cyanide-p-trifluoromethoxyphenylhydrazone (FCCP; the mitochondrial uncoupler), and a mix of antimycin A (inhibitor of Complex III) and rotenone (inhibitor of Complex I) to a final concentration of 1.5 μM, 1 μM, and 0.5 μM, respectively. Subsequently, the oxygen consumption rate (OCR) was measured in time. ATP-linked, basal, maximal, and non-mitochondrial respiration and proton leak were calculated based on the obtained OCR results. The OCR was normalized to the protein concentration determined by the Bradford method.

### Mitochondria and actin staining

Immunofluorescent staining of mitochondrial TOM20 protein and actin filaments was performed on cells plated in 96-well plates (cat. no. 655986, Greiner BIO-ONE, Kremsmünster, Austria) at 0.01 × 10^6^ cells per well. Cells were fixed in 4% formaldehyde in phosphate-buffered saline (Maga-Herba, Legionowo, Poland) for 15 minutes at room temperature, rinsed with PBS, permeabilized with 0.1% Triton X-100 (cat. no. T8787, Merck, Germany) for 10 minutes, rinsed with PBS, and incubated for 30 minutes in blocking PAD solution (bovine serum albumin (BSA, 1% v/v, cat. no. A1595, Merck, Germany) and standard goat serum solution in PBS (10% v/v, cat. no. G9023, Merck, Germany). Cells were incubated with mouse monoclonal primary anti-TOM20 antibody (cat. no. sc-17764, Santa Cruz Biotechnology, Dallas, TX, United States, 1:50) for one hour, followed by 30 minutes incubation with Alexa Fluor 594-conjugated goat-anti-mouse secondary antibody (cat. no. A-11005, Jackson Immuno, Cambridgeshire, United Kingdom, 1:600) and 45 minutes incubation with Phalloidin-Atto 488 conjugate (cat. no. 49409, Merck, Germany). Cell nuclei were counterstained by DAPI (cat. no. MBD0015, Merck, Germany). Cell nuclei were counterstained by DAPI (MBD0015, Merck, Germany). Images were taken and analysed using Axio Observer 7 inverted fluorescence microscope (Carl Zeiss Inc., Dresden, Germany) and ZEN software v.3.3 blue edition (Carl Zeiss Inc., Dresden, Germany).

### Calcium fluctuation assay

The Fluo-4 Calcium Imaging Kit (cat. no. F14217, Invitrogen, Waltham, MA, USA) was used according to the manufacturer’s instructions. Briefly, the culture medium was removed after the treatment and the cells were washed in HBSS + 20 mM HEPES and incubated in a completely humid atmosphere at 37 °C and 5% CO_2_, for 20 minutes, followed by 20 minutes at room temperature with Fluo-4, AM Loading Solution. After the incubation, the cells were washed once in HBSS + 20 mM HEPES and live-cell imagined in HL-1 culture medium using Axio Observer 7 inverted fluorescence microscope with XL PALM-4 S incubator (PeCon GmbH, Erbach, Germany) in a completely humid atmosphere at 37 °C and 5% CO_2_. The pictures of calcium fluctuations were collected using ZEN software v.3.3 blue edition and analyzed using ImageJ FIJI software^31^.

### Statistical analysis

Statistical analyses were performed using GraphPad Prism 9.0 (San Diego, CA, USA). Outliers were identified and excluded using the Robust Regression and Outlier Removal (ROUT) method (Q = 5%). Data distribution was assessed using the Shapiro-Wilk test. However, given the limited sample sizes in some experiments, the results of normality testing were interpreted with caution and supported by visual inspection of data distribution. Depending on the experimental design and data structure, group comparisons were performed using one-way or two-way ANOVA followed by Holm-Sidak, Tukey’s, or Dunn’s post hoc tests, as appropriate. For each experiment, the exact value of *n* is reported. All planned statistical comparisons were performed. The non-annotated comparisons in the figures correspond to non-significant results. Statistical significance was defined as *p* ≤ 0.05, and data are presented as mean ± standard error of the mean (SEM).

## RESULTS

### Irradiated HL-1 mouse cardiomyocytes

In HL-1 cardiomyocytes, irradiation induced divergent metabolic and structural effects. Interestingly, while intracellular ATP concentrations remained stable across both irradiation doses, a differential impact on nucleotide balance and mitochondrial function was observed **(Fig. 1)**. Exposure to the lower dose (10 Gy) disrupted the intracellular energy balance, as reflected by an elevation in intracellular ADP levels and a significant reduction in adenylate energy charge (AEC), suggesting acute bioenergetic stress. This was accompanied by cytoskeletal disorganization, as evidenced by F-actin degradation **(Fig. 2A-B)**, and a decline in mitochondrial respiratory parameters, indicative of impaired oxidative phosphorylation capacity **(Fig. 2C-D)**.

**Figure 1.**
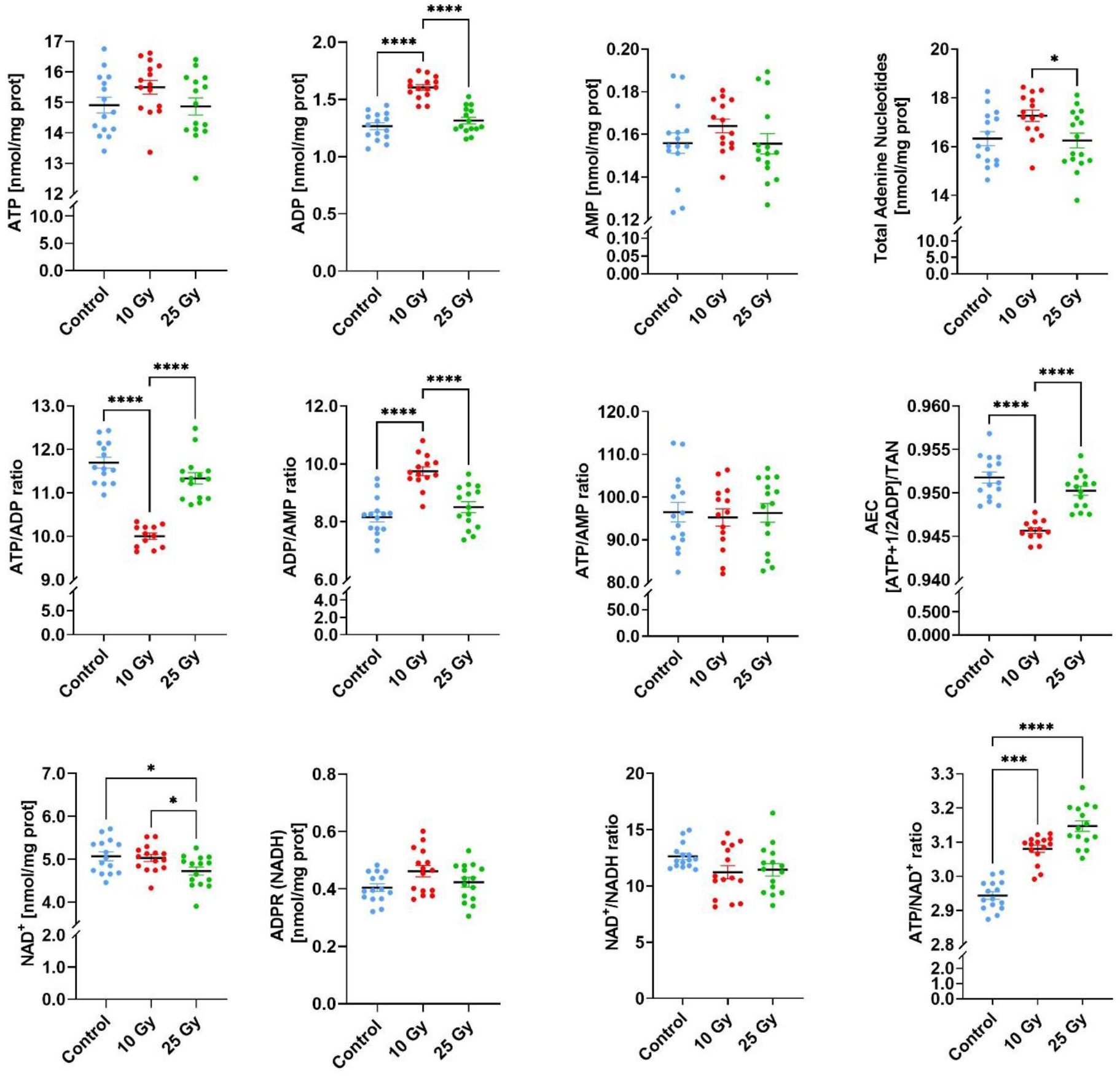
Intracellular nucleotide concentrations in murine cardiomyocyte cell line HL-1 exposed to 10 Gy and 25 Gy doses of radiation. Adenosine-5’-triphosphate **(ATP)**; adenosine-5’-diphosphate **(ADP)**; adenosine-5’-monophosphate **(AMP)**; nicotinamide adenine dinucleotide **(NAD**^**+**^); reduced form of nicotinamide adenine dinucleotide **(NADH)** and their ratios. The energy status of the cells is expressed as an adenylate energy charge **(AEC).** Results are shown as mean ± SEM with individual observations shown as dots, n=15, *p <0.05; ***p <0.001, ****p <0.0001.

**Figure 2.**
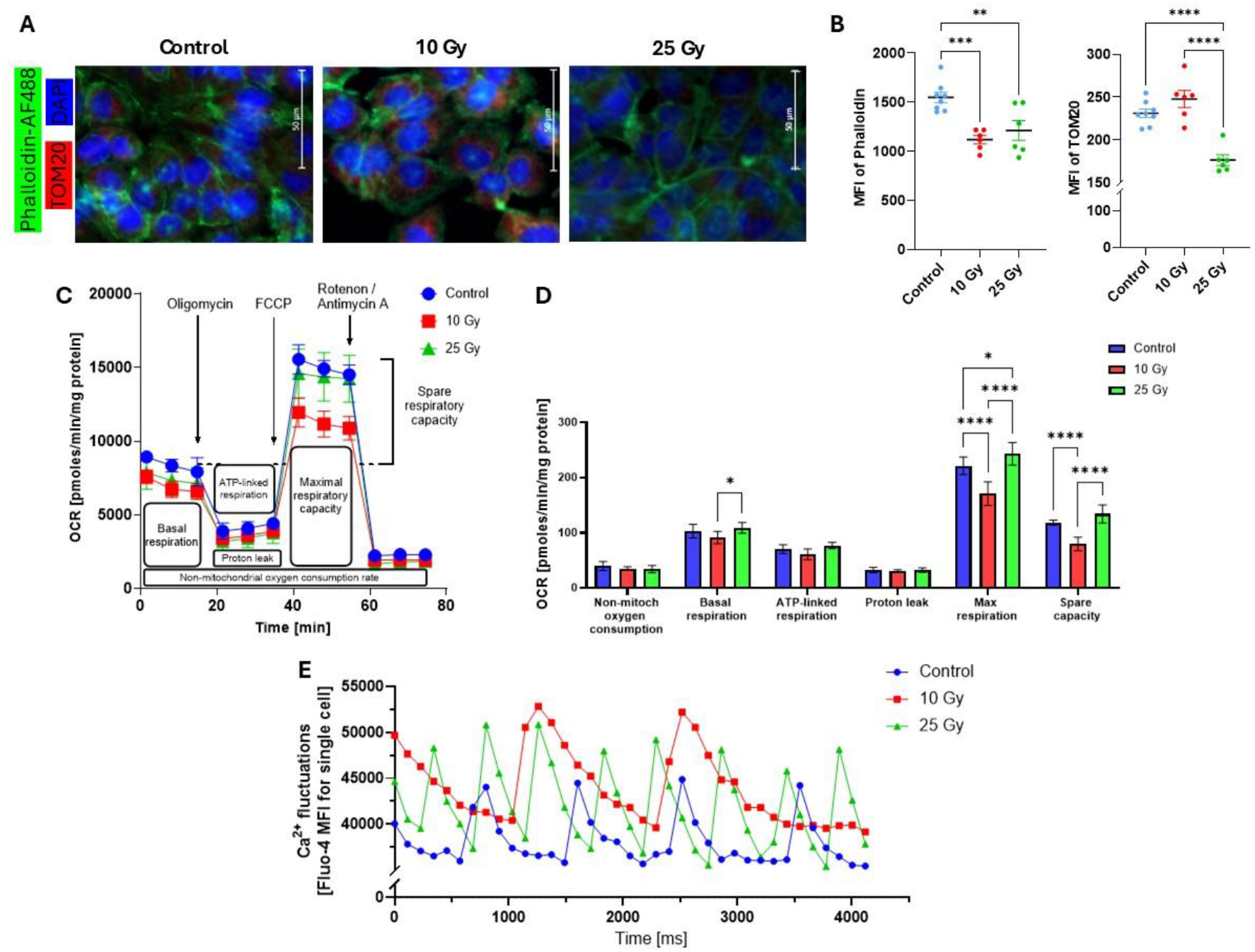
Irradiated cardiomyocyte HL-1 cell line. **(A)** Immunofluorescence staining for actin cytoskeleton (green), mitochondrial TOM20 protein (red), and cell nuclei (blue). **(B)** Quantitative analysis of mean fluorescence intensity **(MFI)** for F-actin and TOM20. **(C-D)** Mito Stress test principle and quantitative results for irradiated HL-1 cells. **(E)** Representative data from a single measurement of Ca2^+^ fluctuations in HL-1 cells assessed in real-time by Fluo-4 staining. Results are shown as mean ± SEM, n=5-8, *p<0.05, **p<0.01, ***p<0.001, ****p<0.0001.

In contrast, the higher dose (25 Gy) did not compromise AEC but led to NAD^+^ depletion and a paradoxical enhancement of mitochondrial respiratory capacity, reflected by increased basal and maximal oxygen consumption rates, as well as spare capacity **(Fig. 2C-D)**. This may point to an adaptive mitochondrial response aimed at compensating for redox imbalance and sustaining energy production. Moreover, high-dose irradiation augmented intracellular calcium fluctuations, specifically increasing their amplitude and frequency, which could potentially contribute to altered cardiomyocyte excitability and arrhythmogenic susceptibility **(Fig. 2E)**.

Collectively, these findings suggest that 10 Gy-dose of radiation impairs cytoskeletal integrity and mitochondrial bioenergetics, while exposure to 25 Gy triggers a metabolic adaptation characterized by increased mitochondrial respiration at the expense of NAD^+^ depletion and calcium dysregulation.

### Bioenergetic response of mice myocardial slices to radiation

In contrast to the HL-1 cardiomyocytes, living myocardial slices from mouse hearts exhibited remarkable metabolic resilience to radiation exposure. Neither 10 Gy nor 25 Gy irradiation induced significant alterations in adenine nucleotide concentrations, catabolite levels, or NAD^+^/NADH redox balance. Key indicators of cellular bioenergetic status, including ATP/ADP ratio and adenylate energy charge, remained stable across treatment groups **(Fig. 3)**. Moreover, no changes in PCr/ATP ratio confirm the overall energy balance maintenance with sustained reserves in the form of phosphocreatine level **(Fig. 4)**. The cardiac cells seem to maintain the phosphorylated form of energy at the expense of free creatine. Simultaneously 10 Gy radiation dose altered guanine nucleotide metabolism specifically by the increase in GMP level.

**Figure 3.**
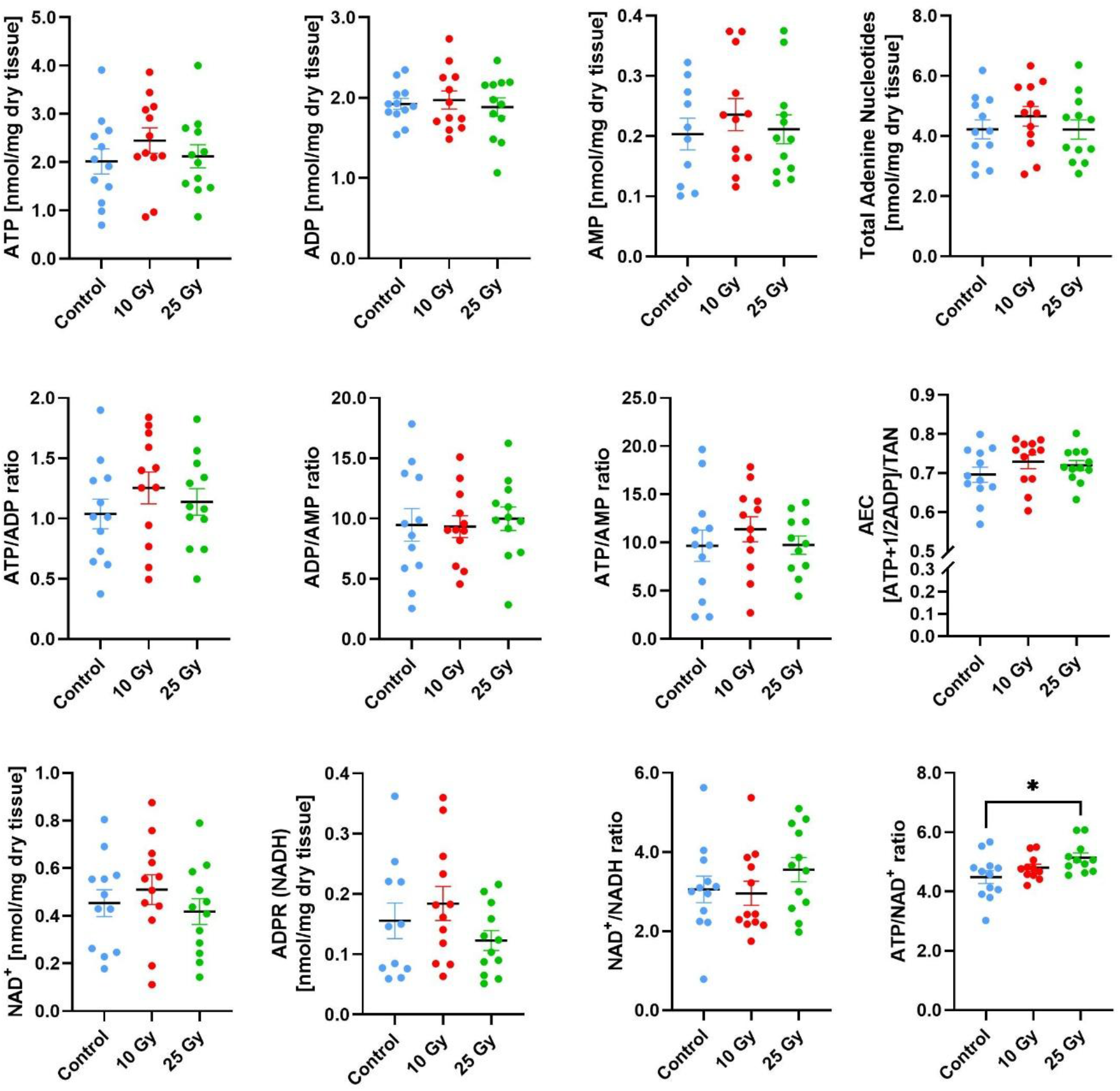
Intracellular nucleotide concentrations in mice left ventricular slices exposed to 10 Gy and 25 Gy doses of radiation. Adenosine-5’-triphosphate **(ATP)**; adenosine-5’-diphosphate **(ADP)**; adenosine-5’-monophosphate **(AMP)**; nicotinamide adenine dinucleotide **(NAD**^**+**^); reduced form of nicotinamide adenine dinucleotide **(NADH)** and their ratios. The energy status of the cells is expressed as an adenylate energy charge **(AEC).** Results are shown as mean ± SEM with individual observations shown as dots, n=12, *p <0.05.

**Figure 4.**
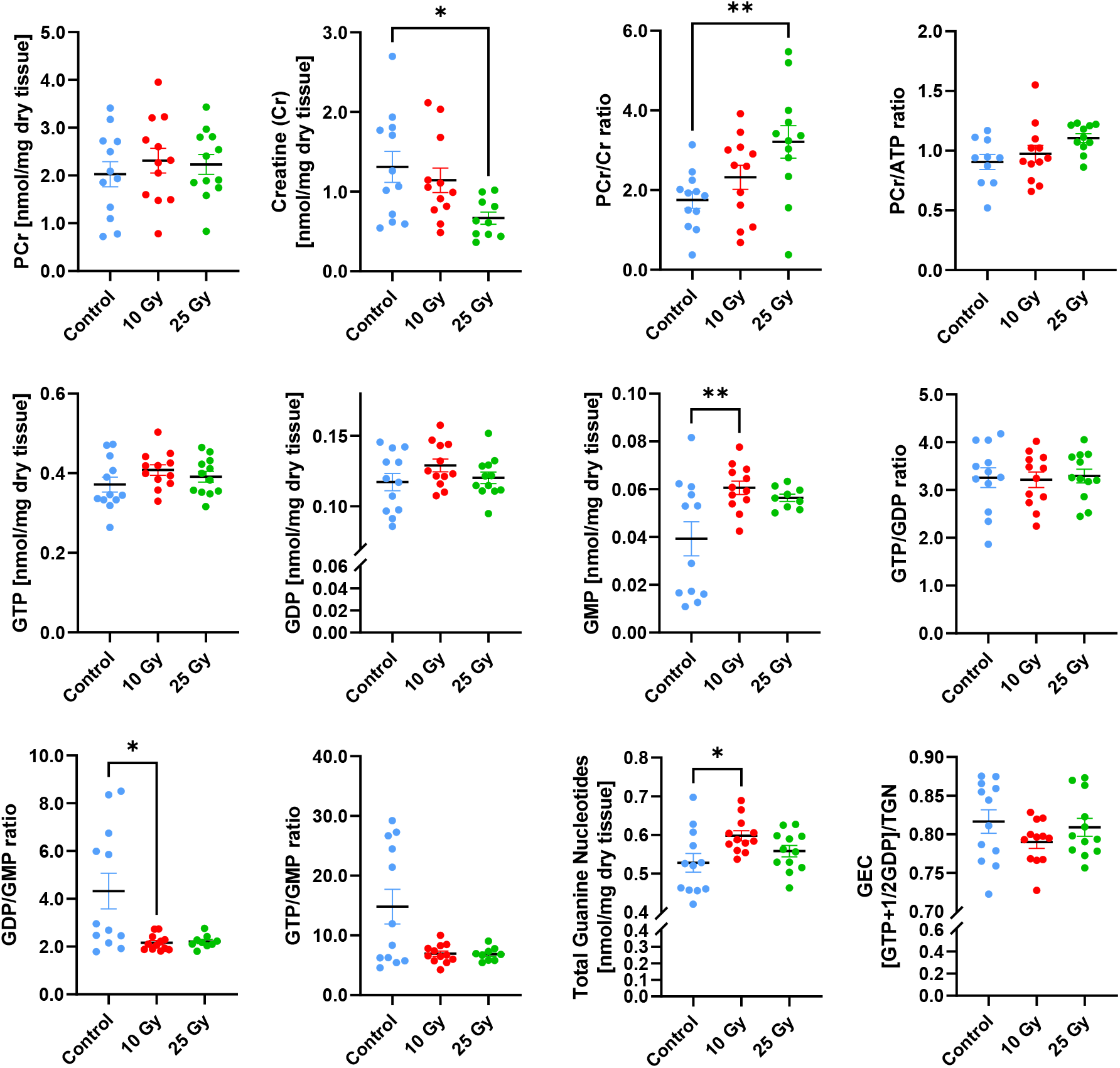
Intracellular guanine nucleotides and creatine/phosphocreatine concentrations in mice left ventricular slices exposed to 10 Gy and 25 Gy radiation doses. Phosphocreatine **(PCr)**; Creatine **(Cr)**; Guanosine-5’-triphosphate **(GTP)**; Guanosine-5’-diphosphate **(GDP)**; Guanosine-5’-monophosphate **(GMP)** and their ratios, Guanylate energy charge **(GEC).** Results are shown as mean ± SEM with individual observations shown as dots, n=12, *p <0.05; **p <0.01.

These findings suggest that, in the intact myocardial tissue context, radiation-induced metabolic disturbances observed in isolated cardiomyocytes are buffered by the multicellular environment and preserved tissue architecture. It is conceivable that paracrine signaling, intact extracellular matrix, or cellular heterogeneity within myocardial slices mitigates the acute metabolic stress induced by irradiation.

### Mitochondrial respiration of irradiated cardiac slices

Irradiation of mouse hearts with 10 Gy or 25 Gy did not significantly alter mitochondrial respiration in cardiac slices compared with non-irradiated controls, as assessed by OCR under baseline conditions after sequential inhibitor injections - oligomycin, FCCP, and a mixture of rotenone and antimycin A **(Fig. 5)**. Non-mitochondrial oxygen consumption was significantly increased in the 25 Gy group compared with both control and 10 Gy slices, whereas basal, maximal, ATP-linked respiration and proton leak showed no statistically significant differences among groups. These findings indicate that high-dose irradiation selectively enhances non-mitochondrial oxygen-consuming processes in cardiac tissue without markedly affecting overall mitochondrial respiratory capacity.

**Figure 5.**
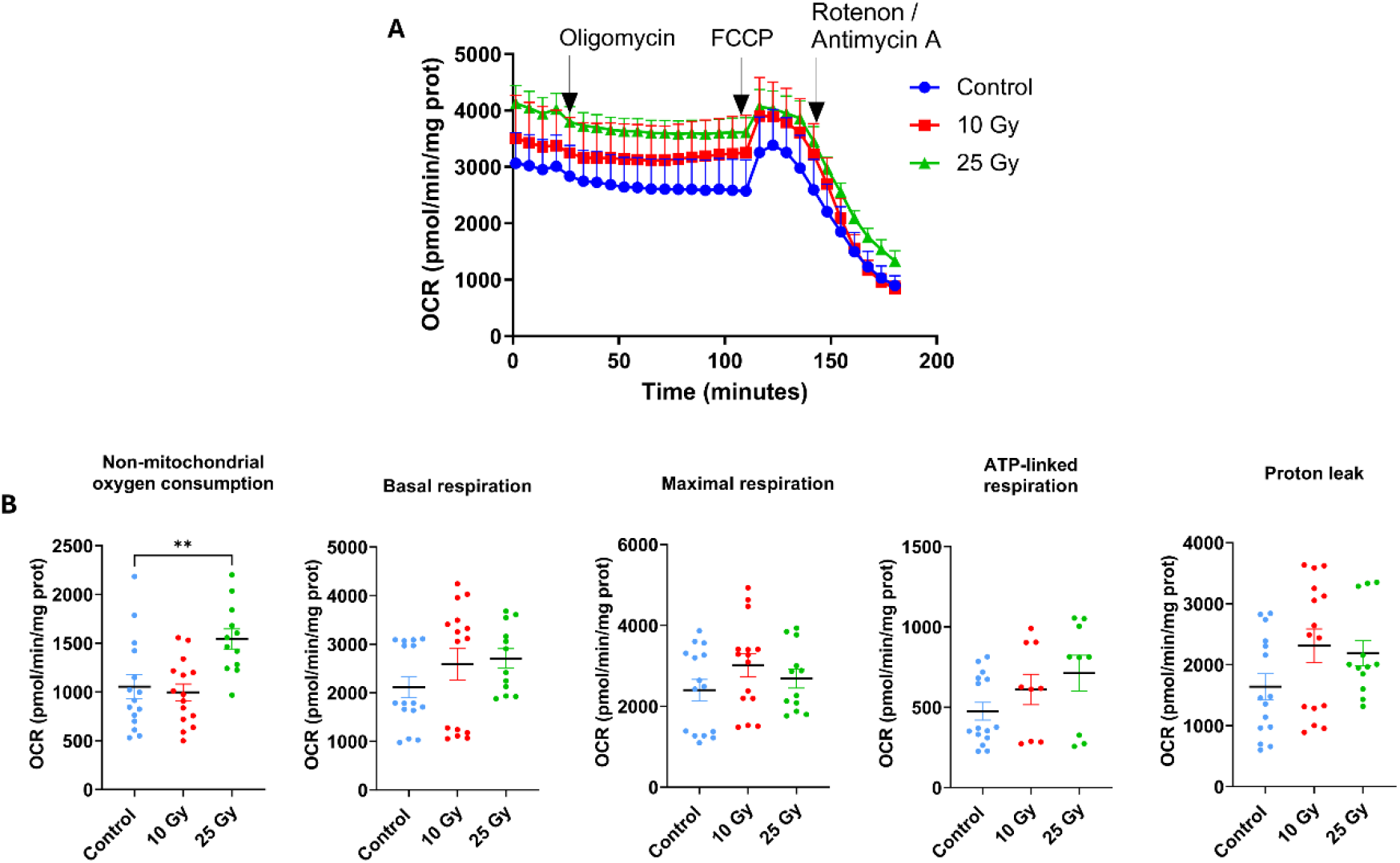
Mitochondrial respiration in ex vivo mouse cardiac tissue slices following 10 and 25 Gy irradiation, presented as (A) Oxygen consumption rate **(OCR)** and (B) quantitative results of basal respiration, maximal respiration, ATP-linked respiration, proton leak, and non-mitochondrial oxygen consumption. Data are presented as mean ± SEM, N=4-5 independent cardiac slices; n=3 biological replicates of oxygen consumption rate per cardiac slice, ** p < 0.01.

## DISCUSSION

In HL-1 cardiomyocytes, we observed a dual, dose-dependent metabolic response to irradiation. Exposure to 10 Gy markedly decreased the adenylate energy charge, ATP/ADP ratio, accompanied by cytoskeletal disorganization, impaired mitochondrial maximal respiration and spare capacity with altered calcium dynamics. These findings indicate a limited ability of mitochondria to increase electron flow in response to energy demand, suggesting partial respiratory chain dysfunction. In contrast, 25 Gy preserved energy charge but induced NAD^+^ depletion and paradoxically increased mitochondrial maximal respiration. This pattern may reflect an acute adaptive response to irradiation-induced redox imbalance. Notably, reduced mitochondrial mass (TOM20 staining) combined with the elevated respiratory parameters suggests activation of compensatory mechanisms aimed at maintaining ATP production despite structural compromise. This adaptive metabolic reprogramming is frequently observed in the early stages of mitochondrial dysfunction. The subsequent rise in ROS levels likely exacerbates oxidative stress, contributing to cellular injury, including cytoskeletal destabilization evidenced by diminished phalloidin staining.

These observations parallel findings reported by Kim et al., who demonstrated increased oxygen consumption rates in irradiated skeletal muscle cells^32^. That metabolic shift was associated with the upregulation of 5′ AMP-activated protein kinase (AMPK), a key regulator of cellular energy homeostasis that enhances oxidative phosphorylation while promoting ATP-generating pathways such as glycolysis and fatty acid oxidation. AMPK activation can occur in response to elevated ADP levels, but also through ROS-dependent mechanisms independent of nucleotide ratios - conditions frequently arising following ionizing radiation exposure^33^.

Conversely, the observed depletion of NAD^+^ after 25 Gy may result from the upregulation of Poly (ADP-ribose) polymerase (PARP) family proteins, which consume NAD^+^ during DNA damage detection, repair, and PAR-dependent cell death pathways^34^. Ionizing directly induces direct single- and double-strand breaks in the DNA, the most potent activators of PARP, whose hyperactivation can drive excessive NAD^+^ consumption and further compromise mitochondrial function.

Our findings also align with recent evidence showing that ionizing radiation triggers functional reprogramming of cardiomyocytes, including alterations in calcium handling and electrophysiological properties ^35^. Disruption of the antioxidant defense balance, especially among superoxide dismutase (SOD), catalase (CAT) and glutathione peroxidase (GPx) has been linked to arrhythmogenic processes. Studies employing 15-25 Gy irradiation demonstrate increased SOD or CAT activities, while GPx activity initially rises as part of an adaptive response and may decline over time, reflecting a multifaceted response of the antioxidant system to elevated oxidative stress^36^.

Calcium dysregulation is a well-established driver of cardiac arrhythmias. Under oxidative stress, mitochondria are highly susceptible to Ca^2+^ overload, which can precipitate irreversible opening of the mitochondrial permeability transition pore (mPTP) and subsequent cell death. Elevated ADP levels have been shown to inhibit mPTP opening, supporting increased mitochondrial Ca^2+^ uptake and preserving mitochondrial integrity, thereby promoting cell survival^37^. However, persistent mitochondrial dysfunction may itself promote arrhythmogenic activity; for example, rapid pacing of cardiomyocytes induces mitochondrial structural alterations and impairs ATP production, contributing to tachyarrhythmogenesis^38^.

Importantly, cardiomyocyte calcium dynamics are tightly coupled to cellular energetic status, including ATP/ADP and NAD^+^/NADH ratios. These metabolic parameters regulate sarco/endoplasmic reticulum Ca^2+^-ATPase (SERCA) activity, ATP-sensitive potassium channels, and ultimately influence action potential duration, Ca^2+^ transient amplitude, and calcium-induced calcium release via RyR_2_ receptors. In tachycardia, elevating NAD levels attenuated SERCA acetylation and facilitated functional recovery of engineered cardiac tissue^39^.

Interestingly, intact murine myocardial slices exhibited remarkable bioenergetic resilience to both 10 and 25 Gy doses of radiation. No significant changes were observed in major nucleotide or redox pools, especially ATP/ADP, NAD^+^/NADH and PCr/ATP ratios. The observed decrease in creatine in the absence of phosphocreatine depletion suggests an alteration in the energy storage equilibrium, potentially reflecting increased creatine utilization or reduced availability, while preserved phosphocreatine levels imply compensatory creatine kinase activity that maintains high-energy buffering capacity. Therefore, it can be assumed that global metabolic balance and mitochondrial function remain stable under the examined conditions.

In our study, 10 Gy irradiation increased the intracellular GMP level, while 25 Gy did not elicit this effect. Cardiac slices also appeared to maintain a high-energy GTP pool at the expense of intermediate metabolites (IMP, hypoxanthine), a pattern consistent with a potential adaptive adjustment of nucleotide metabolism to preserve essential energetic and signaling functions during stress.

When intracellular demand for GTP increases to sustain mitochondrial dynamics^40^, protein synthesis^41^, GTP-dependent signaling^42^, or DNA repair^43^, cells may redirect metabolic flux to ensure sufficient guanylate availability, even at the cost of depleting upstream precursors. Several studies have reported that IMP dehydrogenase activity can increase to preserve guanylate pools^44^, and irradiation-induced metabolic shifts observed here may reflect a similar phenomenon. Moreover, GMP contributes to maintaining the guanylate pool from which GTP is generated, providing the substrate for guanylyl cyclases that synthesize the second messenger cGMP. A post-irradiation increase in GMP could therefore influence cGMP production, with potential downstream effects on vasorelaxation^45^, calcium homeostasis^46^, and broader cellular stress responses.

It is also known that ionizing radiation can modulate the Rho-ROCK signaling axis, affecting endothelial dysfunction, cardiac fibrosis, and structural remodeling^47–49^. Because Rho GTPases require GTP binding for activation, alterations in GMP/GTP availability could, in principle, influence the capacity of cells to engage Rho-dependent signaling. However, such effects are likely to be indirect and context-dependent, and direct causal links between guanylate pool modulation and Rho pathway activation following irradiation remain to be fully established.

Collectively, our findings prove that the multicellular architecture and cell-to-cell interactions preserved in myocardial slices may buffer the acute metabolic stress observed in isolated cardiomyocytes. These observations emphasize the unique translational value of LMS models, which retain the native tissue architecture and multicellular complexity of the heart ^25,50^. The LMS model bridges the gap between simplified *in vitro* systems and *in vivo* animal models. LMS technology preserves the native multicellularity and electrophysiological properties of the myocardium, allowing for high-fidelity assessment of radiation-induced effects on cardiac tissue ^51^. This platform provides an ideal tool for future investigations aimed at identifying therapeutic targets to mitigate radiation-induced cardiotoxicity or enhance the anti-arrhythmic efficacy of SBRT. To understand how SBRT treats ventricular tachycardia at the cellular level, it is important to explore the mechanisms and effects of SBRT under basal non-arrhythmic conditions.

Our findings contribute to the growing body of evidence suggesting that the therapeutic effects of cardiac radiotherapy extend beyond late-onset fibrosis. Previous studies have demonstrated that SBRT exerts rapid anti-arrhythmic effects, often within days of treatment, which cannot be explained solely by structural scar formation. Preclinical data have revealed acute electrophysiological remodeling following irradiation, including increased conduction velocity, reduced action potential duration, and altered gap junction coupling ^52– 54^. Furthermore, a recent clinical and preclinical study demonstrated that cardiac irradiation at relatively low doses (5 Gy) improves cardiac function and attenuates adverse remodeling in murine models of heart failure, potentially *via* suppression of inflammatory and fibrotic pathways ^55^.

Our data extends these observations by providing a detailed analysis of the metabolic footprint of irradiated cardiac tissue. We demonstrate that radiation-induced mitochondrial stress, redox imbalance, and nucleotide depletion are early events that may contribute to both the immediate functional effects and the long-term cardiotoxicity of cardiac radiotherapy. These bioenergetic alterations may underpin the rapid clinical benefit observed after SBRT and offer a plausible mechanistic link to the early electrophysiological changes reported in experimental models.

Importantly, our study addresses a critical gap identified in recent translational research initiatives, including the STOPSTORM consortium, which highlighted the need for mechanistic understanding of radiation-induced cardiac conduction reprogramming ^56^. The significance of this knowledge gap is underscored by ongoing clinical efforts to optimize cardiac radioablation protocols, including both dose escalation and de-escalation trials and expanding indications beyond ventricular tachycardia^57,58^. Interestingly, a recent study described the first patient with refractory VT successfully treated with RT using one fraction of 12 Gy, suggesting that 50% dose reduction might still modify cardiac conduction and result in cessation of refractory VT ^59^. While these initiatives are clinically motivated, they carry the risk of introducing confounding variables and heterogeneity that may limit the interpretability of trial outcomes. Without a clear understanding of the underlying biological mechanisms, it remains challenging to define the optimal dosing strategy and patient selection criteria. Moreover, the absence of mechanistic data may hinder the ability to balance therapeutic efficacy with the potential risk of radiation-induced cardiotoxicity. Our findings contribute to addressing this knowledge gap by providing experimental evidence of early metabolic alterations in cardiac tissue following irradiation. This mechanistic insight is essential to inform rational trial design and to ensure that ongoing clinical efforts are guided by a robust biological framework.

Interestingly, the exposure of mouse cardiac slices to a single high dose of ionizing radiation (25 Gy) does not significantly compromise mitochondrial respiratory capacity in cardiac tissue slices. Basal, maximal, and ATP-linked respiration were preserved, whereas non-mitochondrial oxygen consumption was significantly increased.

This selective increase in non-mitochondrial OCR indicates activation of extramitochondrial ROS-generating pathways, most likely NADPH oxidases, rather than primary dysfunction of the mitochondrial electron transport chain. Consistent with previous reports, high-dose irradiation induces NOX2/NOX4 activation in cardiomyocytes and endothelial cells, promoting superoxide production independent of oxidative phosphorylation. The absence of changes in proton leak further supports preserved mitochondrial membrane integrity at this early time point^60–62^.

The observed metabolic profile suggests that early radiation-induced cardiotoxicity is driven predominantly by redox dysregulation rather than bioenergetic failure. Activation of ceramide-ASMase signaling and mitochondrial-adjacent pathways, including ER-mitochondria crosstalk and calcium handling, may amplify ROS signaling and contribute to subsequent structural remodeling^63^. Together, these findings identify extramitochondrial ROS sources as key early mediators of radiation-induced cardiac injury and potential targets for cardioprotective interventions.

In conclusion, our study reveals that ionizing radiation induces acute and persistent metabolic perturbations in cardiac tissue, characterized by mitochondrial dysfunction, redox imbalance, and nucleotide depletion. These findings provide a mechanistic framework that may explain the early clinical effects of cardiac radioablation and inform future strategies to optimize its efficacy and safety.

## STUDY LIMITATIONS

This study has several limitations. First is the short observation window which precludes definitive conclusions about the long-term metabolic and structural consequences of radiation exposure. The analyses were performed approximately one hour after irradiation, what captures exclusively the early, acute responses and does not reflect the later molecular and phenotypic changes that develop over hours or days. Key processes involved in activation of DNA repair pathways and apoptosis typically emerge over 12-24 hours post-irradiation, therefore the observed alterations do not preclude their occurrence. This temporal limitation constrains the interpretation of functional outcomes and highlights the need for time-resolved profiling in future studies.

Secondly, although our use of LMS model provides a highly translational *ex vivo* model that preserves multicellular architecture and electrophysiological properties, our study lacks *in vivo* validation. Specifically, we did not assess whether the bioenergetic alterations observed in vitro and *ex vivo* also occur in the intact heart of living animals exposed to ionizing radiation. Such validation would require longitudinal metabolic, electrophysiological, and histological assessment in irradiated animal models, providing a functional link between molecular changes and cardiac performance.

Finally, our study focused on mitochondrial function, nucleotide metabolism, and redox balance without addressing other potentially relevant pathways, such as inflammatory signaling, microvascular injury, or autonomic nervous system remodeling, which may contribute to the complex cardiac response to radiation therapy. The interplay between metabolic perturbations, inflammatory responses, and structural remodeling warrants further investigation^12^.

## ACKNOWLEDGEMENTS

B.T. participates in the STOPSTORM Consortium (https://stopstorm.eu/), a publicly funded research network on cardiac radioablation, supported by Horizon 2020 (Grant No. 945119).

## AUTHOR CONTRIBUTIONS

Conceptualization: B.T., B.K.-Z. Methodology: K.S., A.K., A.B., J.K., B.T., B.K.-Z. Formal analysis: K.S., A.K., A.B., K.U., B.T., B.K.-Z. Investigation: K.S., A.K., A.B., J.K., M.P., W.M., K.U., B.T., B.K.-Z. Project administration: B.T., B.K.-Z. Validation: B.T., B.K.-Z. Writing (original draft): K.S., A.K., A.B., B.T., B.K.-Z. (review & editing): K.S., A.K., A.B., J.K., M.P., W.M., K.U., B.T., B.K.-Z.

